# Structures of the human peroxisomal fatty acid transporter ABCD1 in a lipid environment

**DOI:** 10.1101/2021.09.04.458904

**Authors:** Le Thi My Le, James Robert Thompson, Phuoc Xuan Dang, Janarjan Bhandari, Amer Alam

## Abstract

The peroxisomal very long chain fatty acid (VLCFA) transporter ABCD1 is central to cellular fatty acid catabolism and lipid biosynthesis. Its dysfunction underlies toxic cytosolic accumulation of VLCFAs, progressive nervous system demyelination, and neurological impairments including the potentially fatal disease X-linked adrenoleukodystrophy (X-ALD). Molecular details underlying substrate recognition and transport by ABCD1 are poorly understood. Here we determined cryo-EM structures of ABCD1 in phospholipid nanodiscs in a nucleotide bound conformation open to the peroxisomal lumen and an inward facing conformation open to the cytosol at up to 3.5 Å resolution that reveal key details of its transmembrane cavity and ATP dependent conformational transitions. We identify structural elements distinguishing ABCD1 from its closest homologs and show that coenzyme A (CoA) esters of VLCFAs modulate ABCD1 activity in a species dependent manner. Together, our data support a transport mechanism where only the CoA moieties of VLCFA-CoAs enter the hydrophilic transmembrane cavity while the acyl chains extend out into the surrounding membrane bilayer, help rationalize disease causing mutations, and provide a framework for ABCD1 targeted structure-based drug design.

## Introduction

X-ALD (OMIM#300100) is the most common inherited peroxisomal disorder, with a prevalence of approximately 1/20,000 births. Childhood-onset cerebral adrenoleukodystrophy (CCALD) and adult-onset adrenomyeloneuropathy (AMN) are two main forms of X-ALD (*1*). A feature of X-ALD is a build-up of high levels of VLCFAs containing 24 or more carbons throughout the body, which can cause damage to the nervous system due to progressive demyelination (*2*). While X-ALD presents as a metabolic neurodegenerative disorder, phenotypic variability is high (*3*). Dysfunction of the peroxisomal VLCFA transporter ABCD1/ALDP has been identified as an underlying cause of X-ALD, with over 800 disease causing mutations of the *ABCD1* gene identified (*4-7*).

ABCD1 is a peroxisome-membrane spanning protein that mediates the import of various VLCFAs into the peroxisome in an ATP hydrolysis dependent manner (*8-12*) for peroxisome specific beta-oxidation. Accordingly, ABCD1 functional deficiency impairs the degradation of VLCFAs (*13-16*) and may also alter distributions of phospholipid and lysophospholipid species in different brain regions (*17*). ABCD1 is related to and displays functional overlap with two other peroxisomal transporters: ABCD2, also known as ALD related protein/ALDRP (*18*) and ABCD3, also known as peroxisome membrane protein 70/PMP70 (*19, 20*), with which it shares 62% and 39% sequence identity, respectively. Despite substrate overlap, the three transporters display distinct specificities and only mutations in ABCD1 are associated with X-ALD. A fourth ABCD family member, ABCD4 (24% sequence identity to ABCD1), is a lysosome specific transporter that transports vitamin B12 (*21-23*). Unlike its peroxisomal counterparts as well as canonical ABC exporters it shares a fold with, ABCD4 reportedly functions as an importer, with substrate (Vitamin B12/Cobalamin) suggested to bind the ATP bound ‘outward’ facing conformation open to the lysosomal lumen and its cytoplasmic release occurring from an ‘inward open’ conformation (*24*). All four ABCD proteins are half-transporters expressed as single polypeptides containing a transmembrane domain (TMD) and a nucleotide-binding domain (NBD) that must dimerize to form the functional transporter. While some evidence is reported for the existence of ABCD heterodimers (*11, 25, 26*), functional homodimers appear to be most prevalent *in vivo* (*27, 28*).

Structural insights into ABCD1 are currently limited to those gleaned from homology models based on related transporters (*29*) and, more recently, the cryo-EM structure of nucleotide bound ABCD4 in detergent (*24*). To define the ABCD1 structural elements that enable ATP dependent and specific VLCFA transport at the molecular level, we determined it’s structures in nanodisc reconstituted form in both a nucleotide-bound outward open (OO) conformation open to the peroxisomal lumen and an inward open (IO) conformation open to the cytosol. In conjunction with ATP-driven functional assessments, our data reveal the conformational landscape associated with the ATP dependent transport cycle of ABCD1 and highlight key features of its TMD that offer insights into its potential substrate binding mechanism. These structures also allow us to pinpoint the location of the most frequently occurring disease causing mutations in the ABCD1 TMD that will allow for the conceptualization of structure-function hypotheses based on X-ALD patient mutations (*30*). Finally, these structures open the door for more accurate structure guided design of ABCD1 targeted small molecule therapeutics and computational studies of ABCD1 structure and function.

## Results

### ABCD1 *in vitro* activity is modulated by VLCFA-CoAs in a species dependent manner

We utilized a tetracycline-inducible stable cell line to produce human ABCD1 in human embryonic kidney (HEK) 293 cells and tested its ATPase activity in detergent and in liposomes and nanodiscs comprising a mixture of porcine brain polar lipids (BPL) and cholesterol previously used to characterize several other human ABC transporters (*31, 32*) (Fig. 1A, Supplementary Figure 1A). Consistent with previously reported studies (*33*), ATP hydrolysis rates for ABCD1 were in the range of 10 nmol min^-1^ mg^-1^ in liposomes (not accounting for transporter orientation distribution) and nanodiscs, similar to values reported by other groups (*33, 34*), and followed Michaelis Menten kinetics with Michaelis constant (K_M_) values of 0.3-0.4 mM ATP (Fig. 1A). Mutation of a catalytic glutamate residue to glutamine in the Walker B motif in the ABCD1 NBD (ABCD1_EQ_) reduced ATPase rates by ∼50%, similar to observations for ABCD4 (*24*), and ATP hydrolysis was sensitive to inhibition by sodium orthovanadate (VO_4_) or the non-hydrolysable ATP analog ATP*γ*S (Supplementary Figure 1B). We tested CoA esters of various VLCFAs at up to 1 mM concentration for their ability to modulate the activity of ABCD1 as a readout for relevant substrate interactions of in the absence of a direct transport assay. Detergent purified protein was employed to avoid any complications arising from VLCFA-CoA mediated membrane disruption in nanodiscs or liposomal samples. As shown in Fig. 1B, maximal ATPase rate stimulation was observed for C24:0-CoA, followed by C26:0-CoA. Conversely, both C22:6-CoA and acetyl-CoA alone had no observable effect on ATPase activity. To test whether absence of stimulation in the presence of acetyl-CoA could be attributed to lack of a physical interaction with ABCD1, we tested the effect of acetyl-CoA and C24:0-CoA added together. Interestingly, this led to a marked decrease in the stimulatory effect of C24:0-CoA alone, suggesting competition from acetyl-CoA for the same binding site. The implications of these findings in context of substrate transport are discussed below.

**Figure 1:**
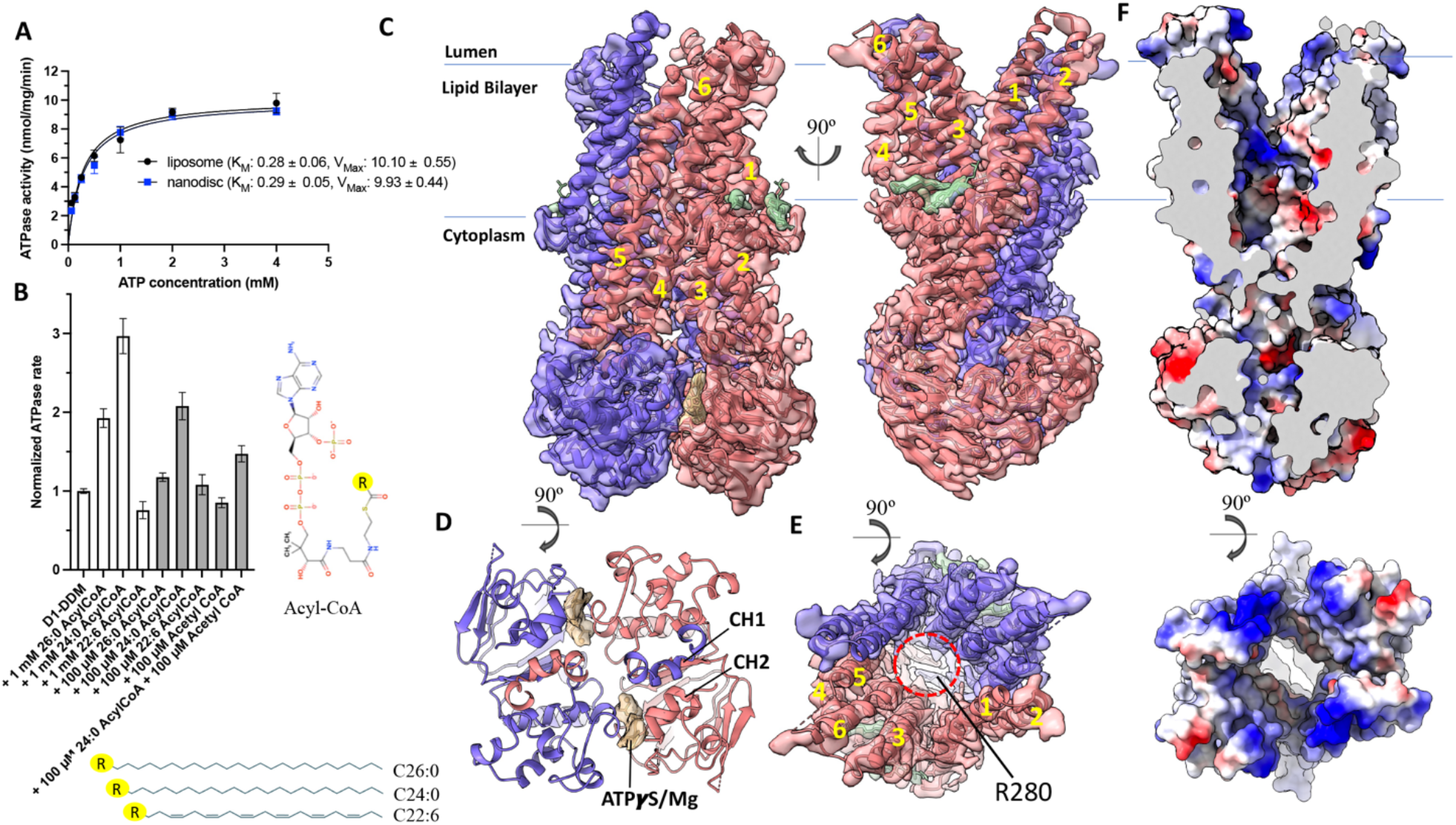
Structural and Functional characterization of ABCD1 human ABCD1. **(A)** ATPase activity of ABCD1 reconstituted in liposomes or nanodiscs as a function of ATP concentration, n=3 and error bars represent standard deviation **(B)** normalized ATPase activity of detergent purified ABCD1 in the presence of CoA esters of varying acyl chain composition, n=6 (100µM) or 9 (1mM) and error bars represent standard error of measurement (**C-E)** Structure of human ABCD1 homodimer viewed from the membrane plane (D), peroxisomal lumen with TMDs removed for clarity, and (E) peroxisomal lumen with intracellular gate circled (dashed red line). EM density (0.4 threshold) and ribbons are colored red and slate for each ABCD1 monomer. EM Density (0.4 threshold) for modeled lipid-like entities and bound nucleotides is colored green and yellow, respectively. TMs are numbered **(F)** Electrostatic surface representation of ABCD1 viewed as a slice from the membrane plane (top) or from peroxisomal lumen (bottom).

### Overall structure of nucleotide bound ABCD1

To determine the structure of human ABCD1 in a lipid environment mimicking its physiological state, we reconstituted detergent purified ABCD1 in BPL/cholesterol nanodiscs utilizing membrane scaffold protein (MSP) 1D1 in the presence of the non-hydrolysable ATP analog, ATP*γ*S (Supplementary Figure 1B). We obtained multiple structures from a single cryo-EM dataset, with a nucleotide bound OO state and an IO state resolved to 3.5 and 4.4 Å, respectively (Fig. 1C, Figs. Supplementary Figure 1D and Supplementary Figure 2). While we observed a range of IO conformations with differing inter-NBD distances, we focus here on the highest resolution of these. The quality of EM density for the higher resolution OO conformation allowed for *de novo* model building of a near complete model of ABCD1. Nucleotide bound ABCD1 adopts a characteristic exporter fold first identified in the bacterial ABC exporter Sav1866 (*35*), entailing a domain swapped architecture where transmembrane helix (TM) 4 and TM5, along with the intervening cytoplasmic helix (CH) from each protomer making extensive contacts with the TMD and NBD of the opposite one (Fig. 1C). Each TMD contains six transmembrane helices that extend well beyond the membrane lipid bilayer in both directions. Two nucleotides, modeled as ATP*γ*S, and two Mg^2+^ ions are bound to the Walker A, Q-loop and ABC signature motifs that exist within the interface between the two NBD (Fig. 1D). The OO conformation is characterized by a large cavity open to the peroxisome lumen and inner peroxisomal membrane. This cavity is lined by polar and charged residues from all 6 transmembrane helices from each protomer and is more hydrophilic and deeper than the corresponding cavities of ABCD4 or Sav1866 (Supplementary Figure 3). It is also considerably wider, especially towards the opening to the peroxisomal lumen where TM5 and TM6 are splayed outward with a noticeable bend in TM6 and display considerable conformational disorder as judged by discontinuous EM density. At the other end of the cavity, a cytoplasmic gate formed by residue R280, stabilized by intrasubunit and intersubunit electrostatic interactions with surrounding residues, seals off access to the cytoplasm (Fig. 1E). The estimated cavity volume of ∼28000 Å^3^ is large enough to accommodate phospholipids present in the surrounding nanodisc. However, no evidence of lipids was found inside the cavity, consistent with its overall hydrophilic nature. Interestingly, we observed lipid-like density features at the outer peroxisomal membrane opening of the OO structure that we tentatively modeled either as cholesterol molecules or unspecified acyl chains (Fig. 1C, 1E). The opening to the inner peroxisomal membrane in the OO cavity also revealed a similar, albeit weaker, stretch of density consistent with an acyl chain. As discussed below, we speculate that the acyl chains of VLCFA-CoAs could occupy similar positions while the CoA moieties bind within the TMD cavity during the substrate transport cycle.

### Transition from OO to IO conformations

Although of lower resolution than that of the OO conformation, the EM map quality of the IO structure allowed for accurate placement of all TMs. Density for the TMD region was of overall higher quality than that for the NBD (Supplementary Figure 2), which were modeled by rigid body placement of the equivalent NBD from the OO structure without modeled nucleotides. The reduced resolution for the NBDs is likely due to increased conformational heterogeneity and is supported by the observation of several 3D classes leading to lower resolution IO structures with varying extents of NBD separation (Supplementary Figure 1D). The final IO model comprised residues 67-443 and 460-684 and allowed for a direct comparison of the IO and OO structures from the same dataset. We observed greater continuity for TM5 and TM6 and the intervening extracellular region (Fig. 2A). This region’s EM density was modeled as a short helical stretch in agreement with secondary structure prediction. Note that this helical stretch is comprised primarily of charged residues that are missing in ABCD2 and ABCD3.

**Figure 2:**
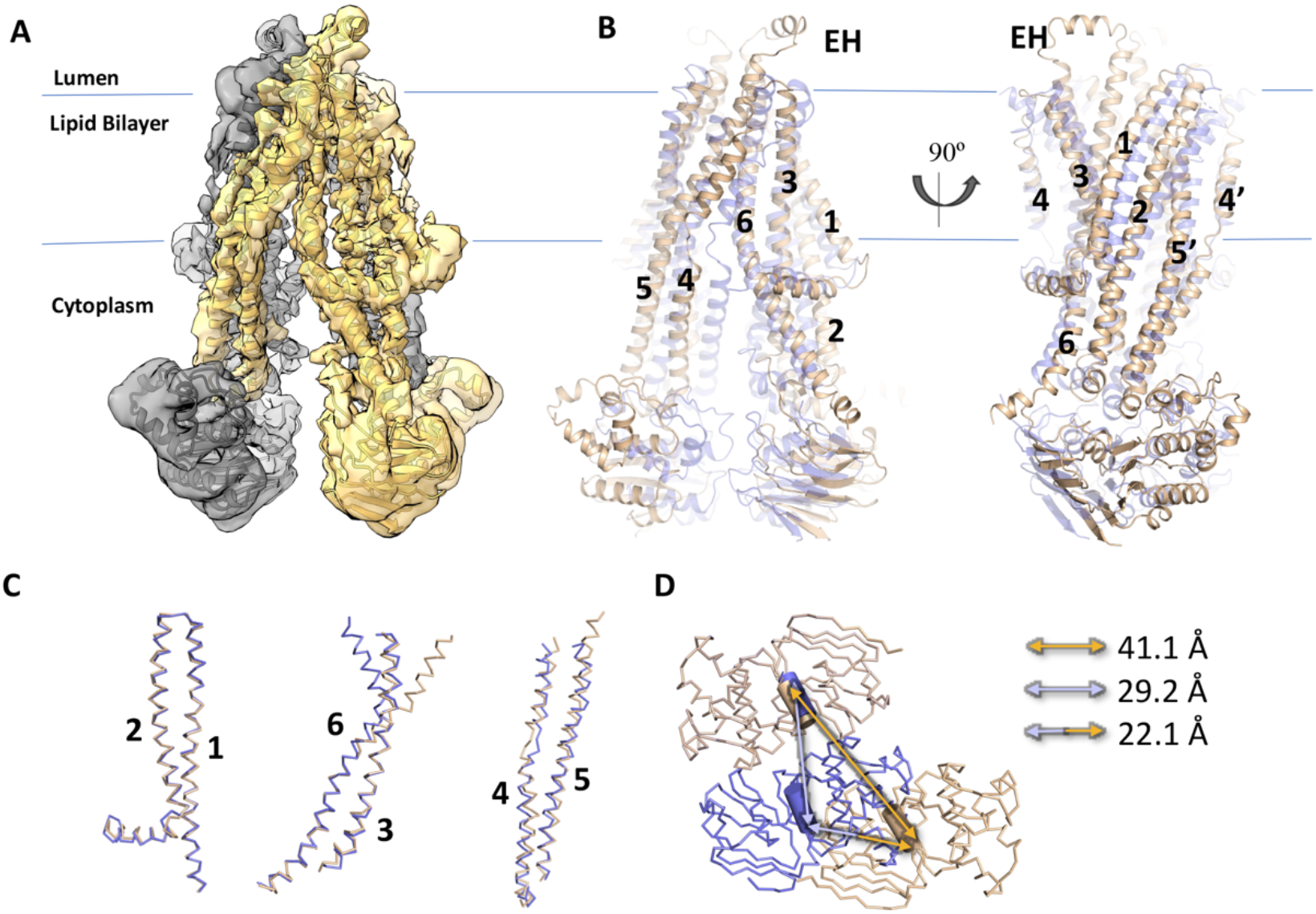
Comparison of IO and OO conformations of ABCD1. **A)** EM density map (0.5 threshold) of human ABCD1 IO (cytoplasm facing) conformation colored yellow and grey for two monomers with similarly colored ribbon representing the protein backbone. **(B)** Overlay of IO and OO structures of human ABCD1 (gold and blue ribbons, respectively) with TMs numbered and external helix (EH) shown for IO conformation. (**C)** Superposition of individual TM pairs from IO and OO structures. **(D)** Comparison of NBDs of IO and OO structures aligned using a single NBD as a reference point. Arrows depict inter coupling helix (cylinders) distances.

Despite the large-scale overall conformational change, TM1-TM2, TM3-TM6, and TM4-TM5 pairs from the IO and OO conformations effectively move as rigid bodies, maintaining their overall conformation, except for a noticeable bend in TM6 in the OO conformation. These transitions follow conserved patterns previously described for type IV ABC transporters/type II exporters (*36*). TM4 in both conformations contains a helical break around residues P263 and G266. G266 is conserved in all three peroxisomal ABCD transporters but not in ABCD4. Helical unwinding of TM4 was also found in ABCB1 and is related to the formation of an occluded conformation with bound substrates or inhibitors in association with its alternating-access mechanism (*31*). A similar break in TM4 of the ABC transporter YbtPQ was proposed as key for substrate release (*37*). The OO-IO transition involves an ∼12 Å translation of the NBDs measured using inter CH1 distances with residue Y296 serving as a reference (Fig. 2D). The resulting conformation creates a large opening to the cytosol, disassembly of the cytoplasmic gating network via repositioning of the residues involved in electrostatic stabilization of R280, and closure of the external cavity and accompanying formation of an external gate that seals the cavity from the peroxisomal lumen (Fig. 3A-C). Like the cavity seen in the OO conformation, the IO cavity is also highly hydrophilic in nature and has openings to the outer peroxisomal membrane leaflet (Fig. 3D), with an estimated volume of ∼37000 Å^3^. As with the OO conformation, no evidence of bound lipids was found within the IO cavity either.

**Figure 3:**
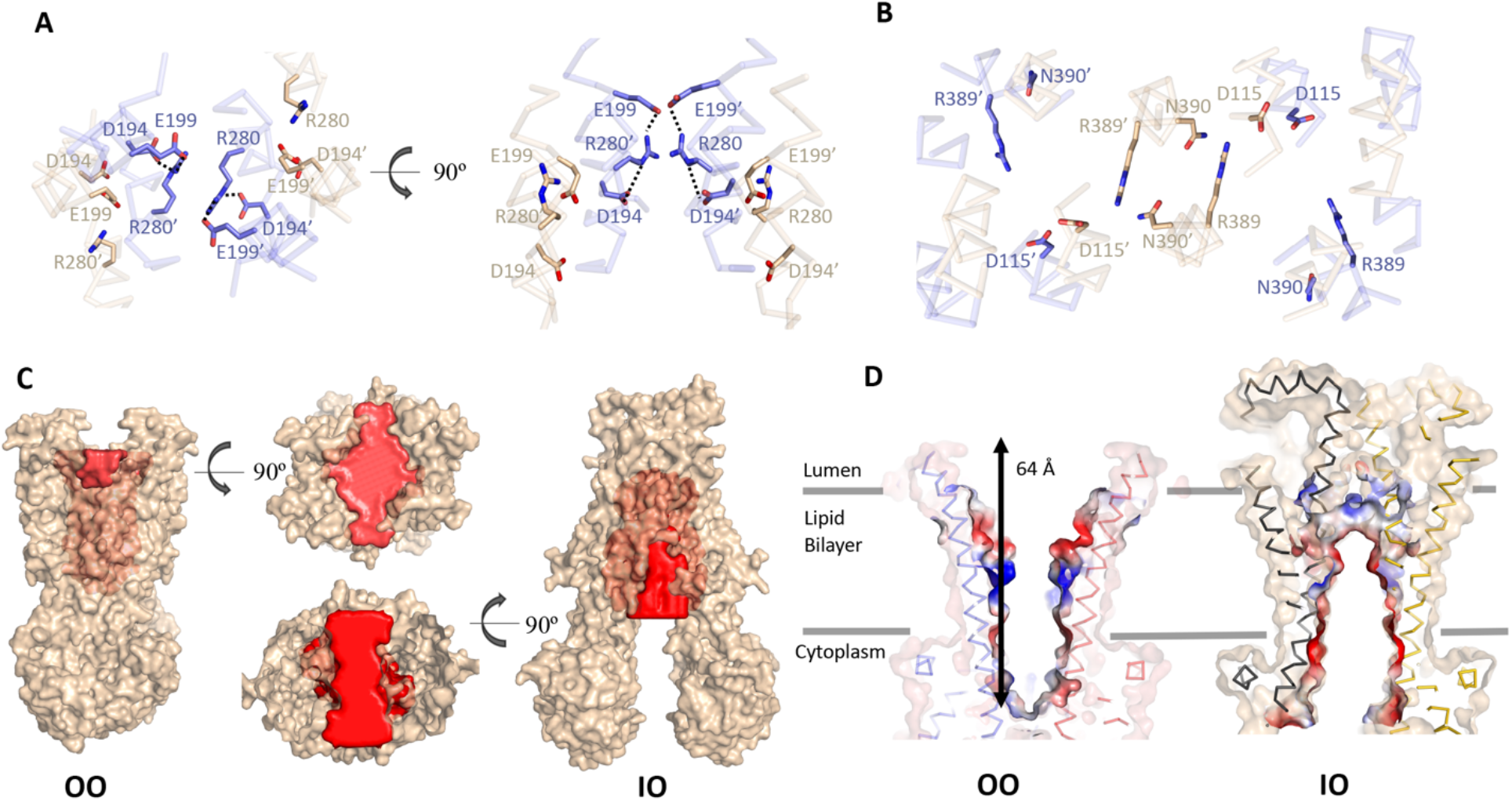
Comparison of IO and OO TMD cavities of ABCD1. **A)** Overlay of IO and OO ABCD1 structures (gold and blue sticks, respectively) but showing intracellular gate and surrounding residues viewed from the peroxisome lumen (left) and membrane plane (right). Equivalent residues from the two halves are distinguished by a prime symbol **B)** Same as (A) showing residues comprising the peroxisomal gate. **C)** Surface representation of OO and IO structures of ABCD1 with solvent exposed cavities colored red. **D)** Slice through electrostatic surfaces of OO and IO structures of ABCD1 focused on the TMD region.

### Analysis of ABCD1 disease causing mutations

To date, over 800 mutations in the ABCD1 gene have been associated with X-ALD. Our ABCD1 structures allow us to pinpoint the location of several single amino acid substitutions and offer a basis for their associated pathogenic phenotype. Due to the high degree of sequence and structural conservation amongst ABC transporter NBDs, we focus here on TMD mutations, which can be further divided into several subsets. The first, a cluster of mutations in TM1 and TM2, occur at the opening to the inner peroxisomal membrane (Fig. 4A). It is plausible that this area is important for substrate transfer through the lateral inner peroxisomal membrane opening of ABCD1 in the OO conformation. A second cluster of mutations is comprised largely of residues from TM4 and TM5 and includes R280 and surrounding residues, which we speculate may lead to destabilization of the OO conformation and disruption of substrate transport. G266, also in TM4, is located at the lateral opening of ABCD1 to the outer peroxisomal membrane in the IO conformation (Fig. 4B). It is one of the most frequently mutated residues in ALDP patients and is likely involved in allowing for secondary structure breaking and kinking of TM4. These data open the door for the design and execution of future *in vitro* mutagenesis studies aimed at analyzing the functional effects of select ABCD1 mutations.

**Figure 4:**
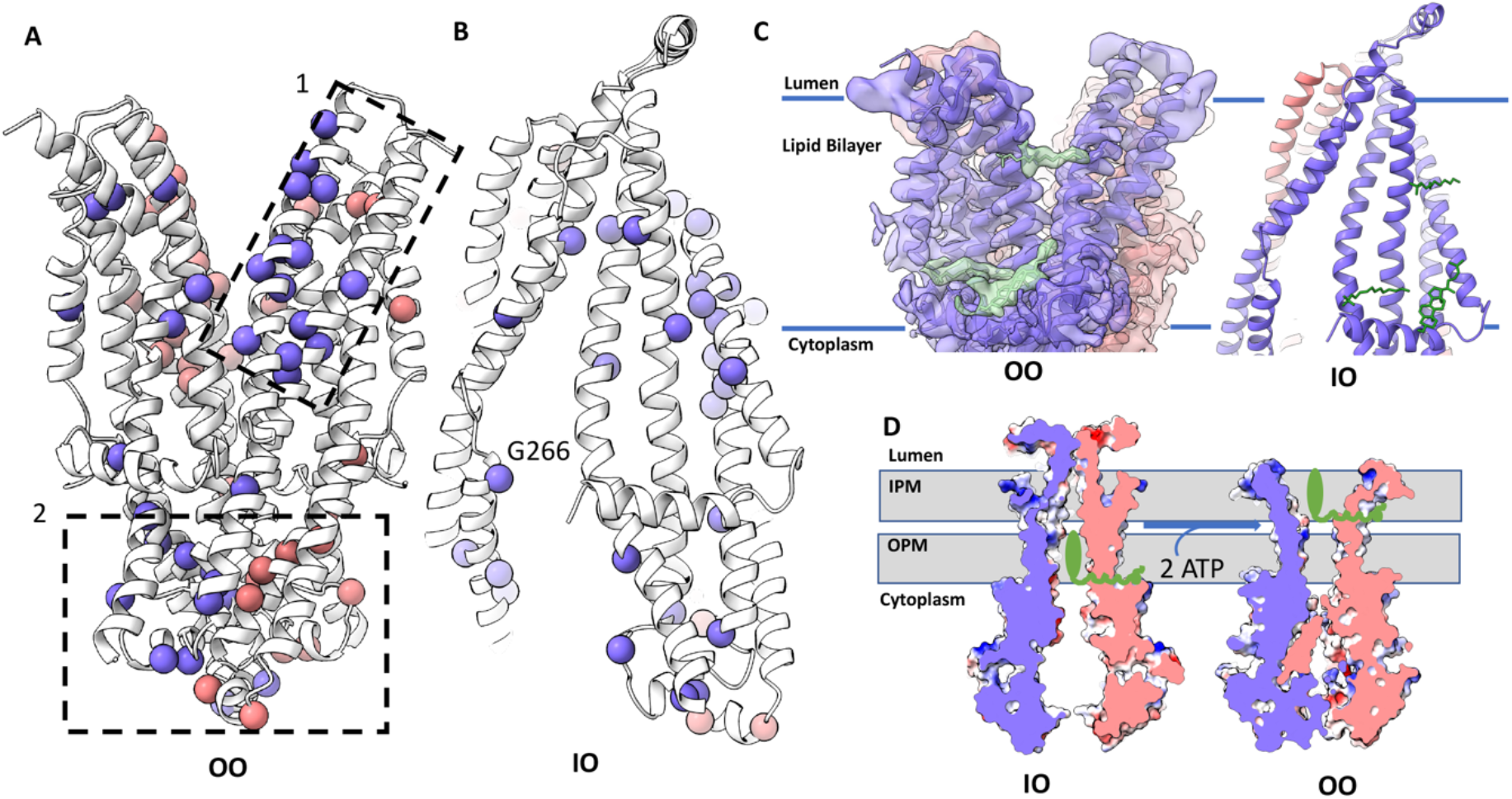
Analysis of TMD mutations in ABCD1 and proposed substrate interactions. **A)** ABCD1 TMD with Cɑ atoms of the most frequently mutated residues shown as spheres colored slate and pink for the two ABCD1 monomers and divided into two main clusters (dashed boxes 1 & 2). **B)** Select ABCD1 disease causing mutations mapped onto ABCD1 IO structure. **C)** EM density (0.4 threshold) of ABCD1 showing modeled lipid-like entities (green) observed in the OO structure (left) and mapped onto the IO structure (right, green sticks). **D)** Hypothetical mechanism of ATP dependent fatty-acyl-CoA (green) translocation across the peroxisomal membrane (grey) mediated by ABCD1.

## Discussion

In the absence of a substrate bound state, the exact mechanisms of substrate recognition and translocation by ACBD1 remain elusive. However, our data offer insights into how VLCFA-CoA may be recognized, with the polar CoA moiety binding within a hydrophilic cavity open to the cytoplasm (IO conformation). The opening to the outer peroxisomal leaflet may offer a portal for the acyl chain to extend outside the cavity into the surrounding membrane space. The ATP dependent switch to the OO conformation would entail a lateral movement of the acyl chain towards the inner peroxisomal membrane. The observation of lipid like density features at the outer peroxisomal membrane opening (Fig. 1C, E, Fig. 4C) is in line with this hypothetical model. When mapped onto the IO ABCD1 structure, the outer membrane density features observed in the OO structure fall at the outer leaflet opening in the IO conformation (Fig. 4C). Moreover, our data show that while acetyl-CoA itself is unable to stimulate ATPase hydrolysis in ABCD1, it can partially inhibit ATPase stimulation by C24:0-CoA. Together, this points to both the binding of the CoA moiety to ABCD1, and the requirement of the acyl chain to trigger the substrate induced conformational changes promoting ATP binding and/or hydrolysis. Acyl chain flexibility may play a key role here, which is supported by our observation that unlike C26:0-CoA and C24:0-CoA, C22:6-CoA has no noticeable effect on ABCD1 ATPase activity.

It is currently unclear whether the fatty acyl chain is separated from its CoA ester for subsequent re-esterification, as has been proposed for the homologous transporter comatose/ABCD1 from *Arabidopsis thaliana (34, 38)*. Our data lay the framework for future studies looking at the exact nature of a substrate bound state of ABCD1, characterization of its thioesterase activity, if any, and, if observed, delineating the mechanism whereby it may be tied to ATP hydrolysis dependent substrate transport. Our structures will also allow for generation of more accurate models for ABCD2 and ABCD3 to shed light on the molecular mechanisms that differentiate these three peroxisomal transporters, which share overlapping but distinct substrate transport profiles. Finally, our structures will prove valuable for design of computational studies aimed at deciphering the fatty acid transport properties and lipid interactions of ABCD1 and provide a solid foundation for structure-based design of correctors or potentiators of ABCD1 for therapeutic use for X-ALD patients.

## Materials and Methods

### Protein expression and purification

We utilized the Flp-In TREX system (Thermo Fisher Scientific) to implement tetracycline inducible expression of human ABCD1 to overcome several impediments associated with weak and inconsistent protein yields from transiently transfected HEK293T cells. Briefly, a synthetic gene construct of isoform 1 of human ABCD1 (Uniprot ID 095477) codon optimized for human cell expression (GeneArt/ Scientific) was first cloned in an expression vector comprising the pXLG gene expression cassette in a pUC57 vector backbone (GenScript) between BamHI and SalI restriction sites as described (*39*). This added an eYFP-Rho1D4 purification tag preceded by a 3C/precision protease site at the C-terminal end of the construct. The full expression construct of ABCD1 or its EQ variant, generated through site directed mutagenesis using forward primer: 5’-CGGCCTAAGTACGCCCTGCTGGACCAGTGTACAAGCGCCGTGTCCATCG-3’ and reverse primer: 5’CGTCGATGGACACGGCGCTTGTACACTGGTCCAGCAGGGCGTACTTAGG-3’ was transferred to a PCDNA5.1/FRT/TO vector between BamH1 and NotI restriction sites and tetracycline inducible stable cell lines were generated as per manufacturer’s protocol. ABCD1 stable cells were grown in Dulbecco’s Modified Eagle Medium (DMEM, Scientific) supplemented with 9% Fetal Bovine Serum (FBS, Gibco), penicillin/streptomycin mixture (Scientific) and antimycotic (Gibco) at 37°C and 5% CO_2_ under humidified conditions. For ABCD1 protein production, cells at a confluency of ∼ 80% were induced with 0.6 µg/ml tetracycline in fresh DMEM supplemented with 2% FBS for an additional 24 hours before being washed with Phosphate Buffered Saline (PBS), harvested and flash frozen in liquid nitrogen. For ABCD1_EQ_ expression, ABCD1_EQ_ stable expressing cells were induced and harvested after 72-hour culture.

For ABCD1/ABCD1_EQ_ purification, thawed cells were resuspended in a lysis buffer (25 mM Hepes pH 7.5, 150 mM sodium chloride (NaCl), 20% glycerol, and 1 mM Dithiothreitol (DTT, Scientific) supplemented with one Complete EDTA free protease inhibitor tablet (Roche) per 50 ml lysis buffer, 800 µM phenylmethylsulfonyl fluoride (PMSF, Sigma), and 20 µg/ml soybean trypsin inhibitor (Sigma). Cell lysate was dounce homogenized (8 strokes with the A pestle) and a 1% Dodecyl maltoside (DDM)/0.2% Cholesteryl hemisuccinate (CHS) (Anatrace) w:w mixture was added for protein solubilization. Protein extraction continued for 90 minutes at 4°C with gentle agitation, followed by centrifugation at 48,000 r.c.f for 30 minutes at 4°C. The supernatant was applied to a Rho-1D4 antibody (University of British Columbia) coupled to cyanogen bromide activated Sepharose 4B resin (Cytiva) and binding was allowed to proceed for 3 hours at 4°C. Beads were subsequently washed with a 4 × 10 bed volume (BVs) of wash buffer (25 mM Hepes pH 7.5, 150 mM NaCl, 20% glycerol, 0.02% DDM/0.004% w:w DDM/CHS, 1 mM DTT). Protein was eluted by incubation with 3 BVs of elution buffer (wash buffer supplemented with 3C protease using 1 milligram 3C per milliliter BV) overnight at 4°C. The his-tagged 3C protease was removed by flowing and washing off cleaved transporter protein over two 100 µl beds of Ni-NTA Superflow resin in tandem (Qiagen). Eluent was concentrated using 100 kDa cut-off Amicon Ultra filters (Millipore-Sigma).

### Nanodisc and liposome reconstitution

Porcine BPL (Avanti) in chloroform were mixed with cholesterol (Sigma) at a final ratio of 80:20 w:w before being dried either under an Argon stream on ice or in a rotary evaporator (Büchi) before resuspension in Di-ethyl ether (Merck) and drying again. For nanodisc incorporation, detergent purified transporter lacking the fusion tag was mixed with membrane scaffold protein (MSP1D1, purified as described (*40*)), and an 80:20 w:w mixture of BPL and cholesterol containing 0.5% DDM/0.1% DDM/CHS using a 1:10:350 (ABCD1:MSP1D1:lipid mixture) molar ratio, and diluted with reconstitution buffer (25 mM Hepes pH 7.5, 150 mM NaCl) to reduce the glycerol concentration to 4% or lower. Nanodisc reconstitution proceeded for 30 minutes at room temperature (RT), followed by detergent removal using 0.8 mg pre-washed Biobeads SM-2 (Bio-Rad) per ml of reaction mix for 2 hours at RT while slowly rolling. ABCD1 or ABCD1_EQ_ nanodiscs washed from Biobeads were concentrated using a 100 kDa Amicon Ultra centrifugal filter (Millipore-Sigma).

For liposome preparation, detergent purified ABCD1 was mixed with a 80:20 w:w BPL/cholesterol lipid mixture at a protein:lipid ratio of 1:10 w:w, following established protocols (*41*) with minor changes. Briefly 0.14% and 0.3% Triton X100 (Sigma) was added to detergent purified ABCD1 concentrated to 1-1.5 mg/ml and BPL/Ch mix, respectively, incubated for 30 minutes at room temperature before being mixed, incubated again for 60 min with gentle agitation. Detergent was removed during successive incubation steps using 20 mg fresh Biobeads SM2 per ml reaction mix each time. The incubation steps were carried out with gentle rolling for 30 min at RT, 60 min @ 4°C, overnight @ 4°C, and 2 × 60 min @ 4°C. The suspension was then centrifuged at 80K rpm for 30 min in an ultracentrifuge. The supernatant was discarded, and the liposomal pellet was washed once with 1 ml of reconstitution buffer per 1 ml original reaction volume. The centrifugation step was repeated, supernatant discarded, and proteoliposomes resuspended in reconstitution buffer to a concentration of 0.5-1 mg^-1^ ml^-1^.

### ATPase assays

Proteins either in 0.02% DDM/0.004% w:w DDM/CHS detergent or reconstituted in 80:20 w:w BPL/cholesterol nanodiscs and liposomes were used in molybdate based calorimetric ATPase assay (*42*) measured at 850 nm on a SYNERGY Neo 2 Multi-Mode microplate reader (BioTek). The assay was carried out at 37°C in a 40-µl volume in 96-well plates. The reaction mixture contained 25 mM Hepes pH 7.5, 150 mM NaCl, 10 mM Magnesium chloride (MgCl_2_), and the protein concentration range was 0.1-0.2 mg ml^-1^. The assays were started by addition of 2 mM ATP, followed by a 40- or 60-minute incubation at 37°C, and stopped by addition of 6% SDS. For ATP K_M_ determination, an ATP concentration range of 0.0625 mM to 8 mM was used. Some experiments included ABCD1 ATPase inhibitors, either 5 mM ATPγS (TOCRIS) or 5 mM orthovanadate and 10 mM MgCl_2_. A higher 5 mM ATP concentration was used in experiments where ABCD1 was incubated with acetyl-coenzyme A (CoA) or very-long-chain fatty acid-CoAs (VLCFA-CoAs: C26:0-CoA, C24:0-CoA, and C22:0-CoA). Two different VLCFA-CoA concentrations (i.e., 1 mM and 100 µM). Statistical analysis was done using GraphPad Prism 9. ATPase rates were determined using linear regression, normalized to the basal ABCD1 ATPase rate. The K_M_ and V_max_ of nanodisc and liposome reconstituted ABCD1 were determined from the fit to the Michaelis-Menten equation of the corresponding ATPase rates. All assays were done at least in triplicate of three independent experimental setups. In assays with substrates, the means included nine replicates and six replicates for experiments with 1 mM and 100 µM VLCFA-CoAs, respectively. All VLCFA-CoAs were purchased from Avanti Polar Lipids and resuspended as per manufacturer instructions. Protein concentrations were measured using gel densitometry analyzed in Image Studio (LI-COR Biosciences) using detergent purified proteins of known concentrations determined by A280 measurements as standards.

### Cryo-electron microscopy grid preparation and data collection

Nanodisc reconstituted ABCD1 was further purified by size exclusion chromatography (SEC) on a G4000swxl SEC column (TOSOH biosciences) pre-equilibrated with reconstitution buffer on an Agilent 1260 Infinity II LC system (Agilent Technologies) at 4°C. Peak fractions were pooled and mixed with 5 mM ATPγS (Sigma) and 10 mM MgCl_2_ for 20 minutes at RT and concentrated to 0.5-1 mg ml^-1^. A volume of 4 µl protein samples was applied to glow discharged Quantifoil R1.2/1.3 grids (Electron Microscopy Sciences, Hatfield, PA, USA) using a Vitrobot Mark IV (Thermo Fisher Scientific) with a 4 sec blotting time and 0 blotting force under >90% humidity at 4°C, then plunge frozen in liquid ethane.

Images were collected under a 300 kV Titan Krios electron microscope equipped with a Falcon 3EC direct electron detector (Thermo Fisher Scientific). Automated data collection was carried out using the EPU software package (Thermo Fisher Scientific) at a nominal magnification of 96,000x, corresponding to a calibrated pixel size of 0.895 Å with a defocus range from -0.8 to -2.6 µm. Image stacks comprising 60 frames were recorded for 60 sec at an estimated dose rate of 1 electron/Å^2^/frame.

### Image processing

All data processing steps were carried out within Relion 3.1 beta and Relion 3.1 (*43, 44*). Motion correction was done using Relion’s implementation of MotionCor2 (*45*) and contrast transfer function (CTF) correction was done using Relion’s gctf (*46*) wrapper. For initial processing, 3,402,440 particles picked from 8378 CTF corrected micrographs extracted at a 3x downscaled pixel size of 2.685 Å/pixel and subjected to multiple rounds of 2D classification. 538,915 particles selected after 2D classification were used for 3D classification (number of classes, K =5) using a rescaled map of the cryo-EM structure of ABCD4 (*24*) (EMDB-9791) as reference. C2 symmetry was applied for all 3D classification and refinement steps. 2 main classes were obtained with NBDs dissociated (IO) or dimerized (OO). The latter was refined to 7.5 Å and used as a reference for 3D classification for the full data set comprising 9040835 particles picked from 19695 image stacks from multiple data collections separated into subset 1 and subset 2. A flow chart depicting the subsequent data processing steps is shown in Supplementary Figure 1D. For subset 1, 2845922 particles after 2D classification were used for 3D classification (K=8) using our initial ABCD1 map as a 3D reference. The 4 highest resolution classes (highlighted blue) were reclassified (K=5) to yield 2 main 3D classes for the OO (highlighted green) and IO (highlighted yellow) conformations. The former was refined to 8.3 Å resolution and the particles from this class were subjected to additional 2D classification from which 257981 selected particles were used for further 3D classification (K=8). The highest resolution of these was combined with an equivalent class from dataset 2 processed similarly to dataset 1. 64750 combined particles were refined to 5.5 Å resolution before being re-extraction using refined coordinates at a pixel size of 0.895 Å / pixel. After 3D refinement and Bayesian polishing, a 3.8 Å resolution map was obtained. Signal subtraction was then done to remove the nanodisc belt before an additional round of 3D classification (K=3). 37237 particles from the highest resolution class was refined using original particles, followed by B-factor sharpening and Post Processing to yield a final map (Map 1) at 3.5 Å resolution. A local resolution filtered map was generated using relion’s own algorithm.

The IO class from dataset 1 was refined to 7 Å resolution before being further 3D classified (K=8). The three highest resolution classes from this round of 3D classification were combined with 2 IO classes from Dataset 2 processed similarly to Dataset 1 and further 3D classified (K=5). 3 highest resolution classes were subjected to 3D refinement and Bayesian polishing to generate a 5.5 Å map. 2 similar classes from another round of 3D classification (K=3) were further refined and the corresponding particles re-extracted to a new pixel size of 1.79 Å/pixel using refined coordinates and subjected to a final round of Bayesian polishing, 3D refinement, B-factor sharpening and Post processing to yield a 4.4 Å map (Map 2) for the IO conformation.

### Model Building

Model building was done in Chimera (*47*) and coot (*48, 49*). Map 1 and its local resolution filtered version were used to build a model of the OO conformation of ABCD1 in coot using a SWISS-MODEL (*50*) generated homology model of ABCD1 based on the published ABCD4 structure as a starting point. The quality of EM density allowed accurate side chain placement for the bulk of the NBD and TMD residues. Restrained real space refinement was carried out in Phenix (*51*). For the IO conformation, TM1-TM2, TM4-TM5, and TM3-TM6 pairs and NBDs from the OO conformation structure were individually rigid body placed into Map 2 followed by manual adjustment as allowed for by the map. The region between residues 361 and 370 was modeled as an alpha helix in agreement with secondary structure predictions. Figures were prepared in Chimera, ChimeraX (*52*), and The PyMOL Molecular Graphics System (Schrödinger, LLC).

## Acknowledgments

We would like to thank the cryo-EM and shared instruments core facilities at the Hormel Institute for help with experimental setup and Dr. Rhoderick Brown, Dr. Jarrod French, and Dr. Jeppe Olsen for critical reading and discussion during manuscript preparation. This work was supported in part by the Hormel Foundation (Institutional research funds to AA), the United Leukodystrophy Foundation (ULF research grant to AA), and the EAGLES Cancer Telethon Postdoctoral Fellowship (to LTML).

## Author contributions

AA conceived the research. LTML, JRT, and AA performed all research with technical assistance from PXD. JB helped with EM sample preparation and handling and data collection. LTML, JRT, and AA wrote the manuscript with input from all other authors.

## Data and materials availability

The cryo-EM Maps for nanodisc reconstituted human ABCD1 have been deposited at the Elecron Microscopy Databank (EMDB) under accession codes EMD-24656 (Map 1, ABCD1-OO) and EMD-24657 (Map 2. ABCD1-IO) respectively. The associated atomic coordinates have been deposited at the Protein Data bank (PDB) under accession codes 7RR9 (ABCD1 OO) and 7RRA (ABCD1 IO).

---

**Supplementary Figure 1:**
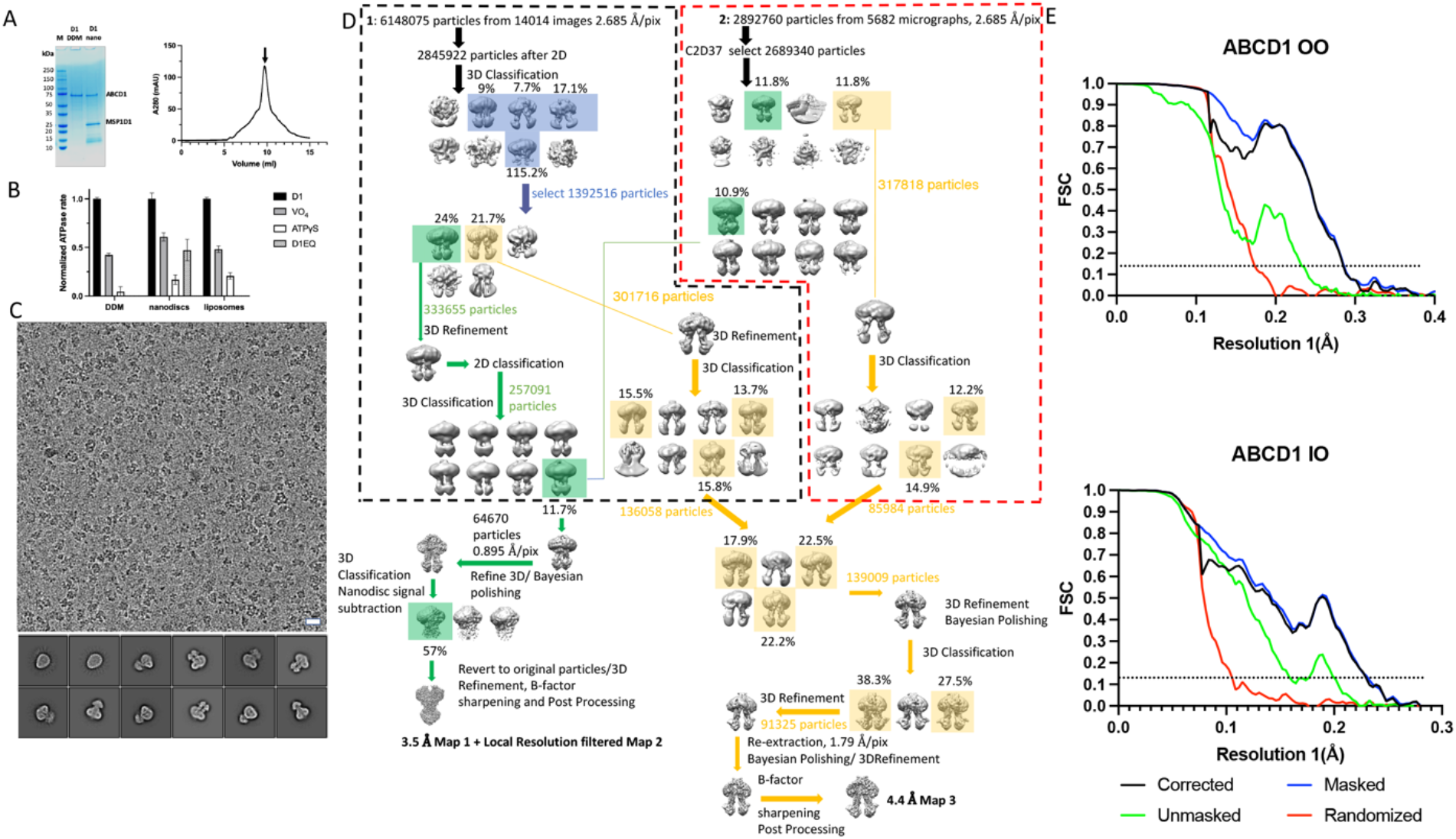
Functional reconstitution and structure determination of nanodisc reconstituted human ABCD1. **A)** 4-15% Coomassie stained gradient gel for detergent purified and nanodisc reconstituted ABCD1. **B)** ATPase activity of wild type ABCD1 (D1) in the presence of sodium orthovanadate (VO4) and ATP*γ*S. ATPase rates in each case are normalized to activity of wildtype ABCD1 alone. N=3, error bars represent S.D **C)** Representative micrograph of nanodisc reconstituted ABCD1 at ∼2.5 µM defocus along with representative 2D classes. **D)** Data processing flowchart. Classes leading to OO and IO structures are highlighted green and yellow, respectively. Dashed lines represent individual datasets before classes were combined for further processing. **E)** Fourier Shell correlation curves for ABCD1 IO and OO maps. Dashed lines indicate 0.143 cutoff.

**Supplementary Figure 2:**
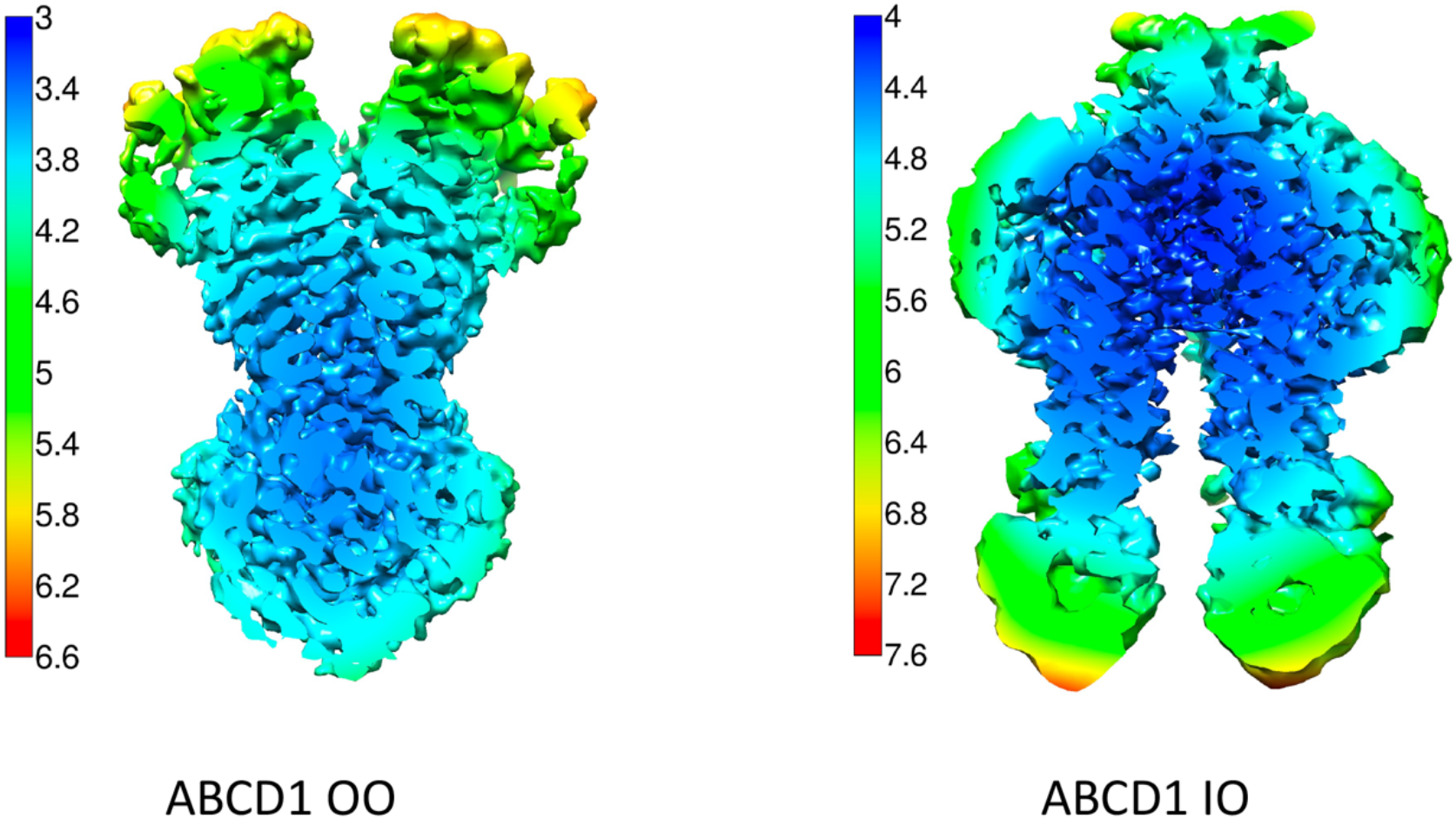
Local resolution variation in ABCD1 OO and IO maps. Central slice through local resolution filtered EM maps of nanodisc reconstituted human ABCD1 OO (left) and IO (right) conformations with color key shown on left of each.

**Supplementary Figure 3:**
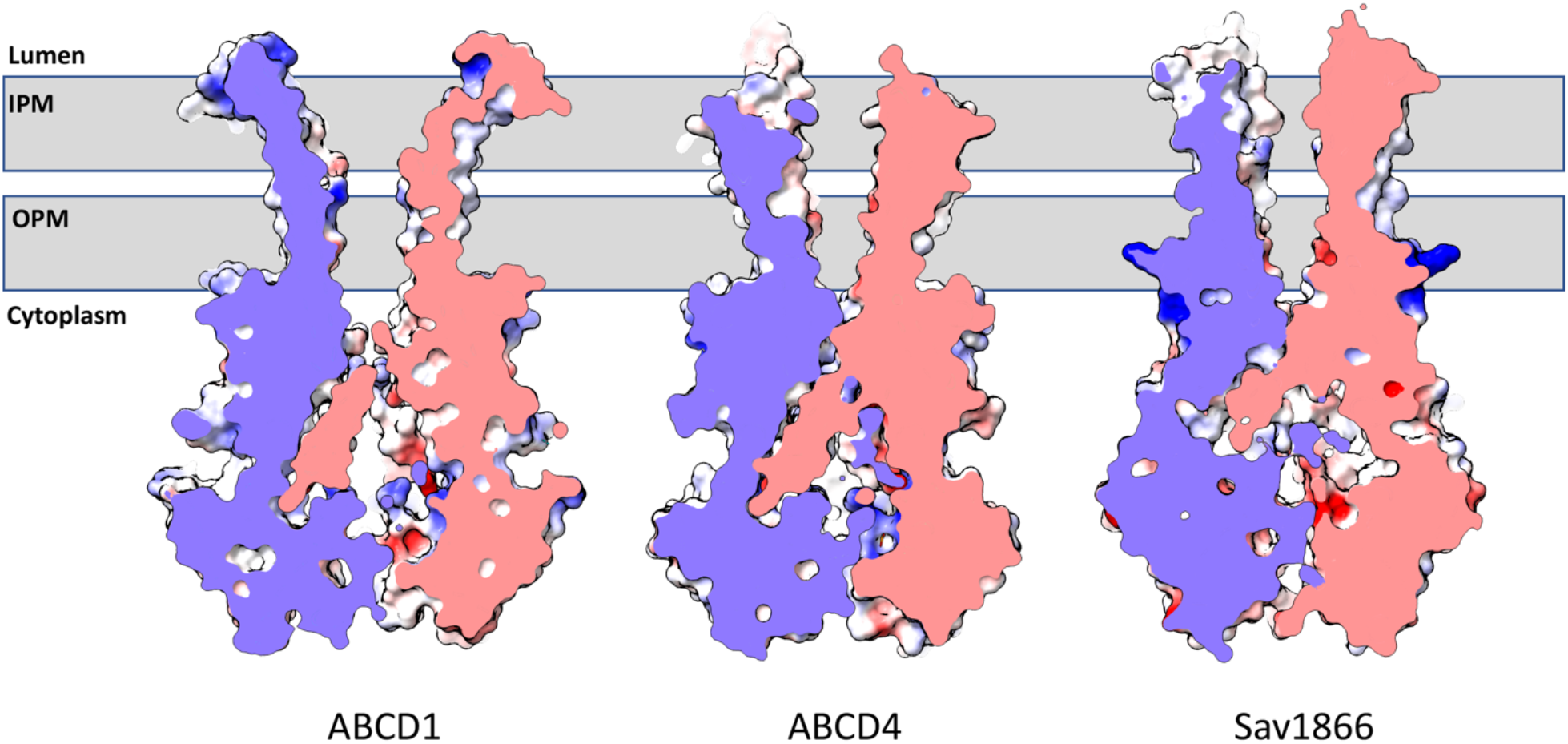
Comparison of ABCD1 transmembrane cavity to that of ABCD4 and Sav1866. Grey bar represents approximate position of peroxisomal membrane. IPM= Inner peroxisomal membrane, OPM=outer peroxisomal membrane.

**Supplementary Figure 4:**
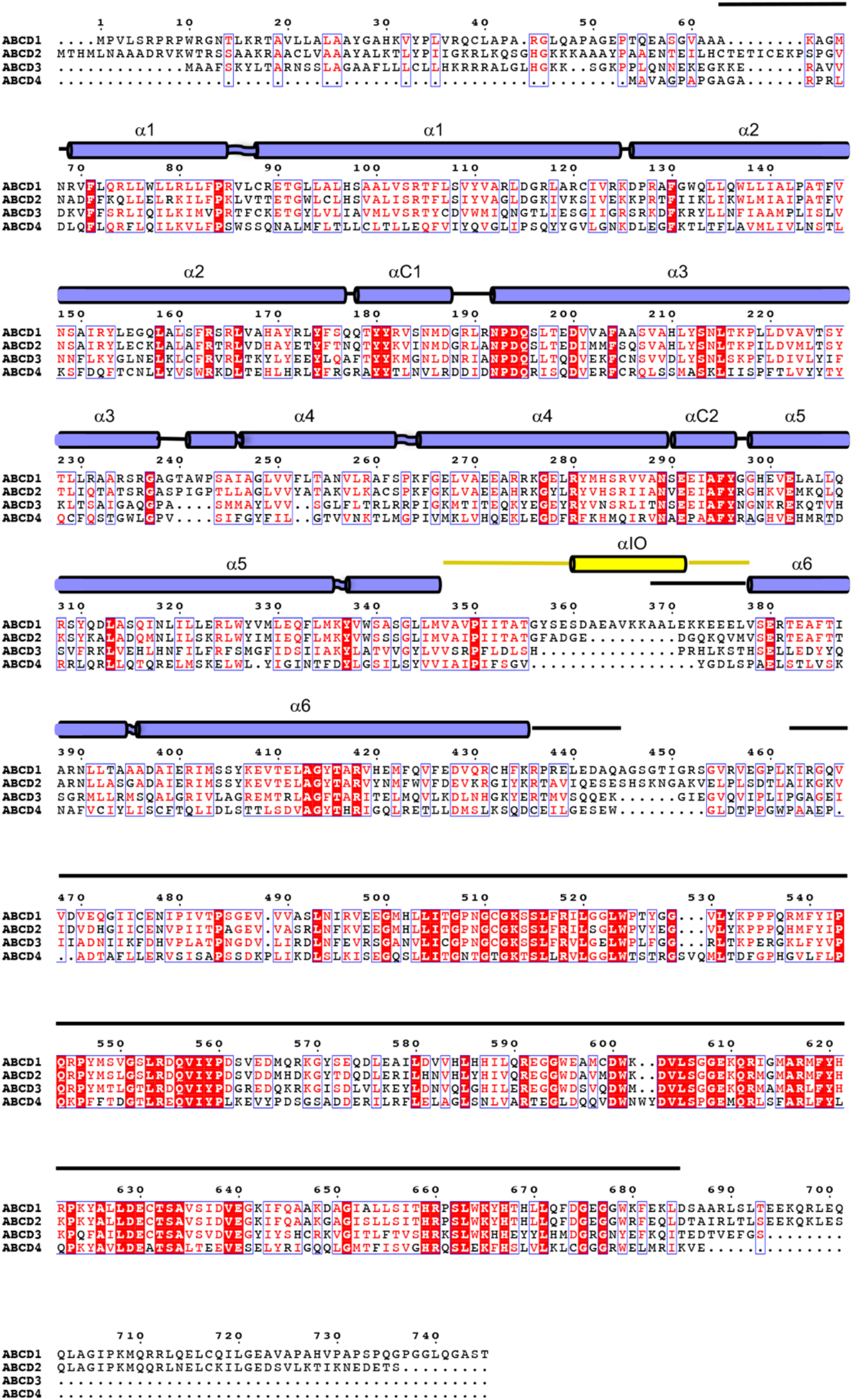
Sequence alignment of human ABCD family transporters with secondary structure assignment based on ABCD1 structures presented in this study shown.

**Supplementary Table 1:**
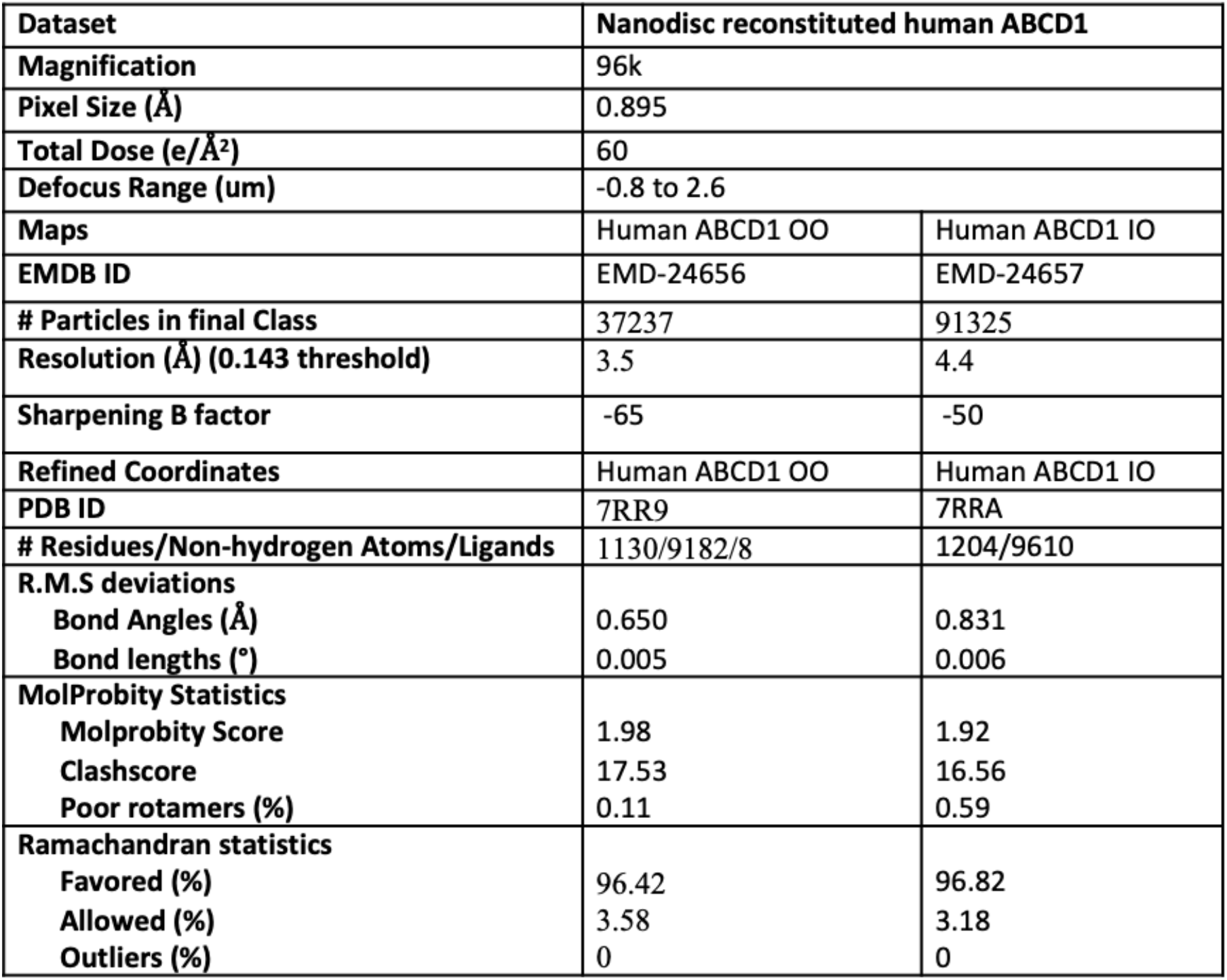
Data processing and refinement statistics.

